# Comparison of miRNA profiling methods using synthetic miRNA pools and standardized exRNA samples reveals substantial performance differences

**DOI:** 10.1101/645762

**Authors:** Paula M. Godoy, Andrea J. Barczak, Peter DeHoff, Srimeenakshi Srinivasan, Saumya Das, David J. Erle, Louise C. Laurent

## Abstract

MicroRNAs (miRNAs) found in biofluids play functional roles in health and in disease pathogenesis, underpinning their potential as clinical biomarkers. Several platforms for measurement of extracellular RNAs have recently become available. We evaluated the reproducibility, accuracy, sensitivity, and specificity of four miRNA quantification platforms, including one widely used discovery approach (small RNA-seq) and three targeted platforms (FirePlex, EdgeSeq, and nCounter). Using pools of synthetic miRNAs, we observed that reproducibility was highest for RNA-seq and EdgeSeq, that all three targeted platforms had lower bias than RNA-seq, and that RNA-seq had the best ability to distinguish between present and absent sequences. Overall reproducibility was lower for plasma samples than synthetic miRNA pools. We compared expression of placental miRNAs in plasma from pregnant and non-pregnant women and observed expected differences with RNA-seq and EdgeSeq, but not FirePlex or nCounter. We conclude that differences in performance among miRNA profiling platforms impact their relative utility as potential assay systems for clinical biomarkers.

## INTRODUCTION

MicroRNAs (miRNAs) are a class of small (18-22 nt) non-coding RNAs with known roles in gene regulation (Bartel, 2004). miRNAs can be released from cells and have been detected in over 15 biological fluids (Godoy et al., 2018, Sohel, 2016) and have been proposed as possible biomarkers for tissue pathologies. Accordingly, there is strong interest in using miRNAs as biomarkers for prediction and diagnosis of disease or for monitoring response to therapy (Das et al., 2019). Several miRNA measurement methods exist and have been used extensively, including quantitative PCR (qPCR), microarrays, and small RNA-sequencing. Previous studies that have systematically compared the performance of these methods have provided a useful resource for those studying miRNA expression (Mestdagh et al., 2014, Giraldez et al., 2018, Yeri et al., 2018). While small RNA-seq is excellent for discovery studies, it is less useful for high-throughput or rapid turnaround applications, which are important factors for validation studies or clinical assays. Recently-developed platforms catered specifically to assay extracellular RNAs (exRNAs) seek to address these gaps. Understanding differences in reproducibility, bias, and ability to detect biological differences across these platforms is important for selection of methods for translation of initial discovery-based exRNA studies to large scale clinical validation and actual clinical use. Here, we aim to build upon previous studies by comparing a previously assessed small RNA-sequencing protocol to three relatively novel platforms: HTG Molecular’s EdgeSeq miRNA Whole Transcriptome Assay (EdgeSeq), Abcam’s FirePlex (FirePlex), and NanoString’s nCounter (nCounter).

The platforms used in the study vary widely. Among the tested methods, small RNA-seq is the only one that does not use probes specific to miRNAs of interest. The small RNA-seq method we use here was optimized for low-input samples and was shown to be less biased than other widely used commercial small RNA-sequencing methods (Giraldez et al., 2018). This was primarily achieved by modifying both the 5’ and 3’ adapters to include 4 randomized nucleotides on their 3’ ends that are ligated to the RNA molecule.

EdgeSeq is a multiplexed nuclease protection assay with next generation sequencing readout (Girard et al., 2016). First, probes containing sequences complementary to specific miRNAs and flanking sequences for downstream amplification are incubated with the miRNA-containing sample. Probes that successfully hybridize to their cognate miRNA in the sample are protected from nuclease digestion, amplified with the addition of barcodes, and then sequenced. Therefore, the output for EdgeSeq is read count, as in small RNA-seq, but unlike small RNA-seq, the number of reads reflect the quantity of probes that were bound by miRNAs and thus protected from digestion.

The remaining two methods use probes and fluorescent reporters. The Multiplex Circulating FirePlex miRNA Assay (Abcam, Cambridge, MA) is based on gel microparticle technology (Chapin et al., 2011). The FirePlex hydrogel particles contain a central region that binds specific miRNAs based on complementarity, as well as two separate end regions with differing fluorescent intensities that serve as a barcode for the central analyte region. Bound miRNAs are ligated to universal adapters, then eluted from the hydrogels for amplification by PCR using biotinylated primers specific for the universal adaptors. Following amplification, the now-biotinylated miRNA targets are rehybridized to the hydrogel particles and a fluorescent reporter specific for biotin is used for quantitative detection of fluorescence on a flow cytometer. Analysis of the fluorescence attributed to the biotin-specific reporter (representative of relative target miRNA abundance) combined with the unique fluorescent barcodes on each end of the hydrogel (specific signature for the miRNA target) allows for multiplexed detection of up to 68 individual miRNAs per assay.

The nCounter platform relies on hybridization of miRNAs to probes conjugated to unique fluorescent barcodes and can potentially assay up to 800 different targets (Denaro et al., 2017). Unlike the other platforms we tested, nCounter does not require an amplification step and counts the total number of fluorescent barcodes to determine the quantity of miRNA molecules in the sample. EdgeSeq and FirePlex can use isolated RNA or crude biofluid as input, while small RNA-seq and nCounter require isolated RNA.

miRNA measurement using each of the tested platforms involves several steps, each of which can display preferences for certain RNA molecules, resulting in differences in the efficiencies with which different miRNAs are detected. For example, during small RNA-seq library preparation, adapters ligate more efficiently to some miRNAs than others (Jayaprakash et al., 2011 and Hafner et al., 2011), whereas in hybridization-based assays, efficiency of probe binding varies in a sequence-specific manner, leading to cross-hybridization (Wu et al., 2005).

Additionally, incorporation of incorrect nucleotides can be introduced during amplification or sequencing, leading to alignment errors or cross-hybridization. These target-specific biases preclude using signal strength as a direct measure of the abundance of a particular miRNA and can lead to differences in the ability to detect and reproducibly quantify specific miRNAs.

The NIH-supported Extracellular RNA Communication Consortium (ERCC) was launched to establish foundational knowledge and technologies for extracellular RNA research (Das et al., 2019). Here we report the results of an ERCC-supported miRNA analysis platform comparison that examined reproducibility, bias, specificity, and relative quantification using both defined pools of synthetic miRNAs and pooled human plasma samples. The use of synthetic miRNA pools allowed us to assess performance using complex mixtures of miRNAs at known concentrations. The use of plasma samples allowed us to compare performance using a highly relevant sample type and assess the ability of two platforms to assay miRNAs directly, without RNA isolation.

## RESULTS

### Reproducibility across technical replicates for synthetic miRNA pools

Three pools of synthetic miRNAs (see Methods) were analyzed with each of the four platforms. The first pool, referred to as the equimolar pool, contained 759 synthetic human miRNAs and 393 synthetic non-human RNA oligonucleotides at the same molar concentration. The other two pools, referred to as ratiometric pools A and B, each contained the same 286 human miRNAs and 48 non-human miRNAs but at different concentrations, with the absolute concentrations of individual miRNAs varying over a 10-fold range within each pool. The relative concentrations of a given miRNA between pool A and pool B varied from 1:10 to 10:1. To assess the reproducibility of each platform, we examined the coefficient of variation (CV) of each miRNA’s signal intensity across technical replicates for RNA-seq, EdgeSeq, and FirePlex (Table S1, Table S2). Only miRNAs considered to be detectable were included in the analysis (see Methods). Technical replicates were not performed by NanoString and therefore reproducibility could not be assessed for the nCounter assay. For the equimolar pool, the median CV was higher for FirePlex (22.4%) than for small RNA-seq (8.2%) and EdgeSeq (6.9%). CV decreased as signal increased for RNA-seq and EdgeSeq but not for FirePlex (Fig. 1). Ratiometric pools A and B showed similar CVs as the equimolar pool, although CVs decreased as signal intensities increased for all platforms, including FirePlex (Supplemental Figure 1, Table S2, Table S3). Overall, we concluded that technical reproducibility was higher for small RNA-seq and EdgeSeq than for FirePlex.

**Figure 1.**
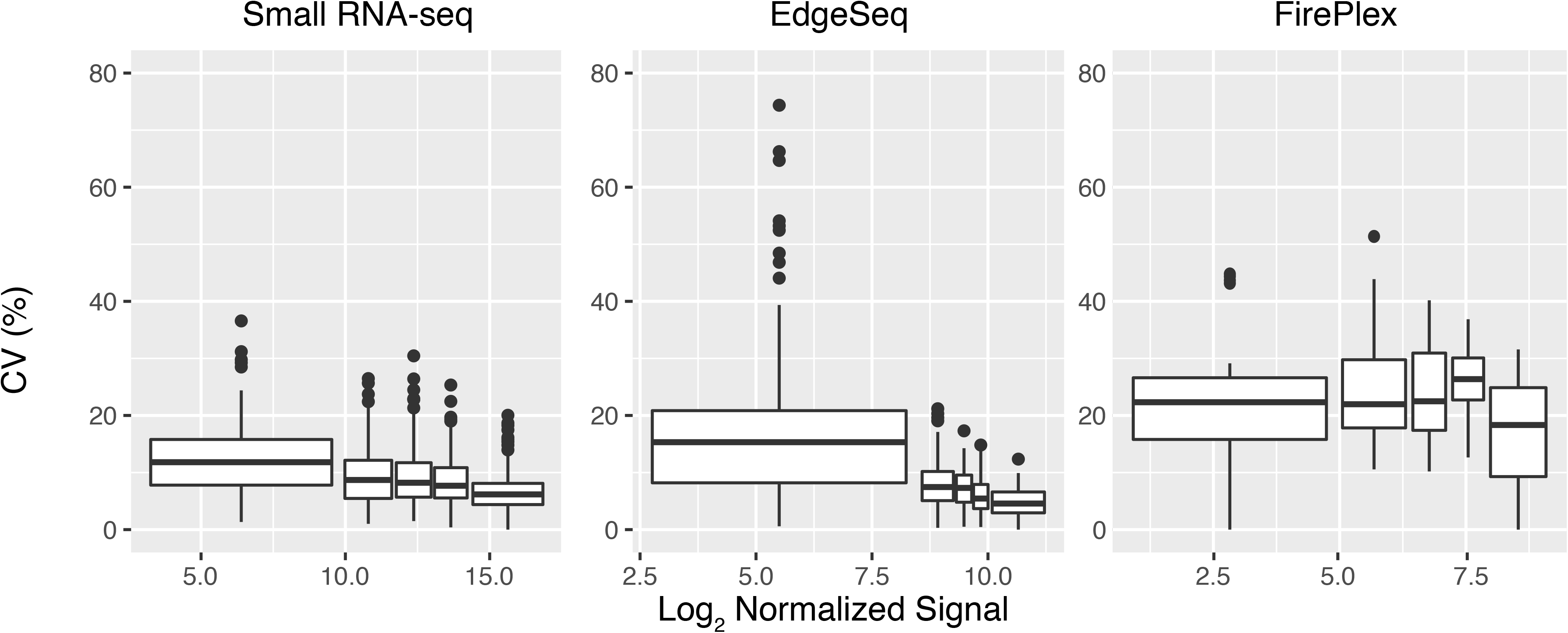
Reproducibility of small RNA-seq, EdgeSeq, and FirePlex when using a Synthetic Equimolar Pool. Coefficient of variation for technical replicates (small RNA-seq N = 4, EdgeSeq N = 3, FirePlex N = 3) expressed as a percentage as a function of median signal. Each boxplot represents ~20% of the total number of detectable miRNAs, grouped by ascending expression (for total number of detectable miRNAs: small RNA-seq N = 988, EdgeSeq N = 451, FirePlex N = 103). Boxes represent median and interquartile ranges, whiskers represent 1.5 times the interquartile range. Dots represent outliers.

### Assessing the bias associated with each platform

Equal quantities of two different miRNAs can result in different signal intensities due to detection bias. Determining detection bias for a set of miRNAs requires a comparison between the amounts of these miRNAs and the signal intensities associated with each miRNA. In biological samples, the miRNA concentrations are usually not known. To accurately assess the bias for each platform we used the equimolar and ratiometric pools. For each pool, we calculated the expected signal intensity based on the known concentration of each component miRNA that is considered detectable and quantified the detection bias as the ratio of observed to expected counts (see Methods).

Small RNA-seq exhibited the most bias, with many target RNA sequences displaying substantially lower than expected signal (log_2_ detection bias < 0) and a relatively small number of target RNA sequences showing markedly higher than expected signal (log_2_ detection bias > 0, Fig. 2). With small RNA-seq, only 31% of the miRNAs in the equimolar pool had signals that were within 2-fold of the median signal. EdgeSeq had the least bias (76% within 2-fold of the median signal), while nCounter (47%) and FirePlex (57%) were intermediate. Results obtained with the ratiometric pools were similar to results obtained with the equimolar pool (Fig. 1). Therefore, although the version of small RNA-seq that we used has lower bias than some other widely used RNA-seq methods, it nonetheless exhibited substantially more bias than the other three platforms.

**Figure 2.**
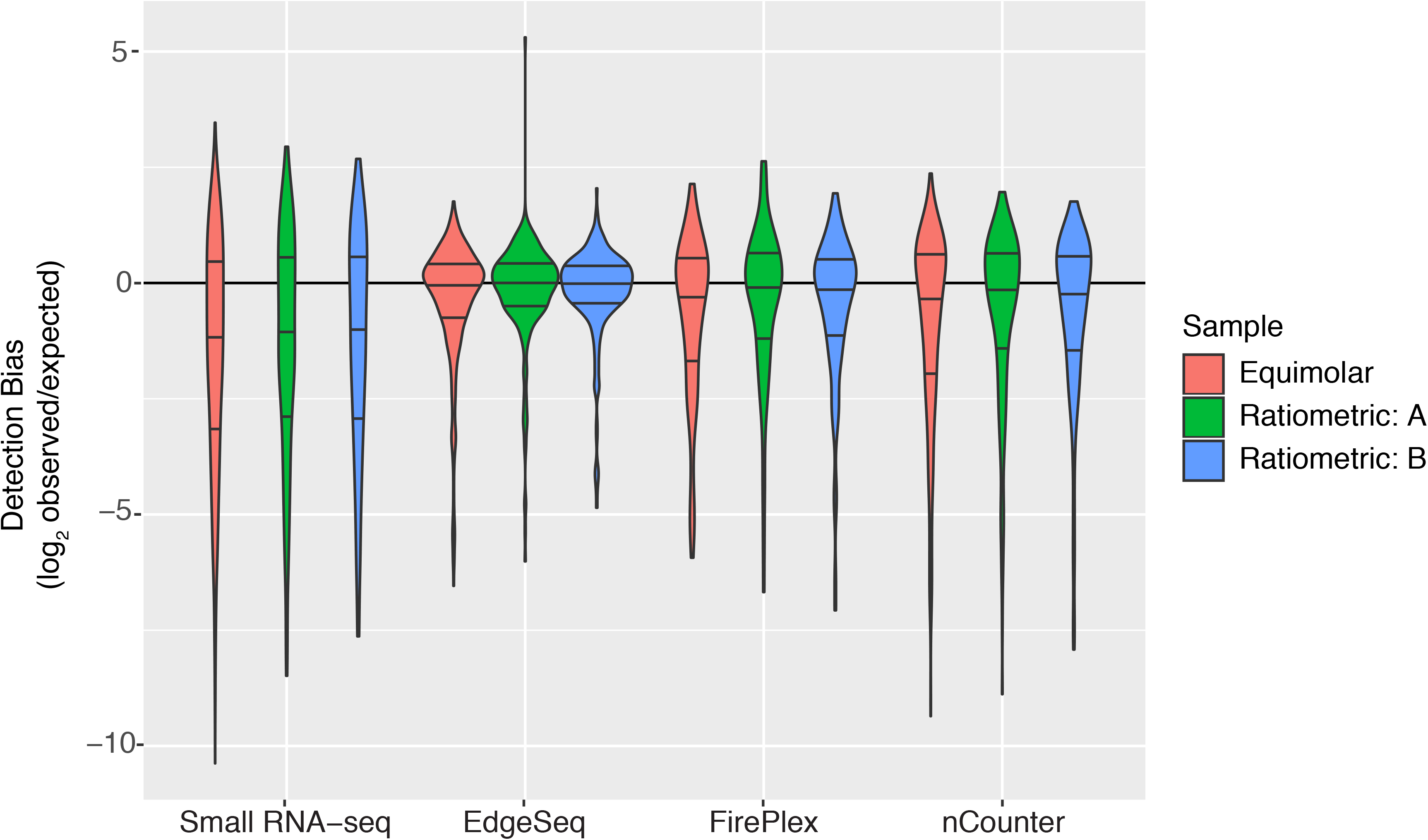
Detection Bias Determined Using Synthetic miRNA Pools. The observed signal to expected signal (detection bias) is plotted for each platform for the equimolar sample (red), the ratiometric A pool (green), and the ratiometric B pool (blue). The width of the violin plot represents the density of miRNAs. The lines represent the median (middle) and interquartile ranges (top and bottom).

Relationships between detection bias and miRNA sequences differed between platforms. For small RNA-Seq, EdgeSeq, and to a lesser extent FirePlex, the GC content of an RNA sequence correlated with the detection bias (Fig. 3). For EdgeSeq, miRNAs that were least efficiently detected all had low GC content (<35%). There was minimal if any evidence of an association between bias and GC content for nCounter (Fig. 3). We also explored associations between the bias and the identity of the 5’ or 3’ nucleotide of the target miRNA for each of the platforms (Supplemental Figure 2). In EdgeSeq, signal intensities of miRNAs with a 5’ cytosine (N = 64) were higher than those with a 5’ uracil (N = 200) (Bonferroni-adjusted p = 1.5 × 10^−3^). Both small RNA-seq and nCounter had higher signals for miRNAs with 3’ guanines (N = 218 and N = 85, respectively) compared to 3’ uracils (N = 365, N = 175, respectively; p = 4.2 × 10^−3^ and p = 1.6 × 10^−2^, respectively). Overall, GC content and the identities of the 5’ and 3’ nucleotides had effects on signal intensity that differed between platforms, but these factors are not sufficient to accurately predict or adjust for bias.

**Figure 3.**
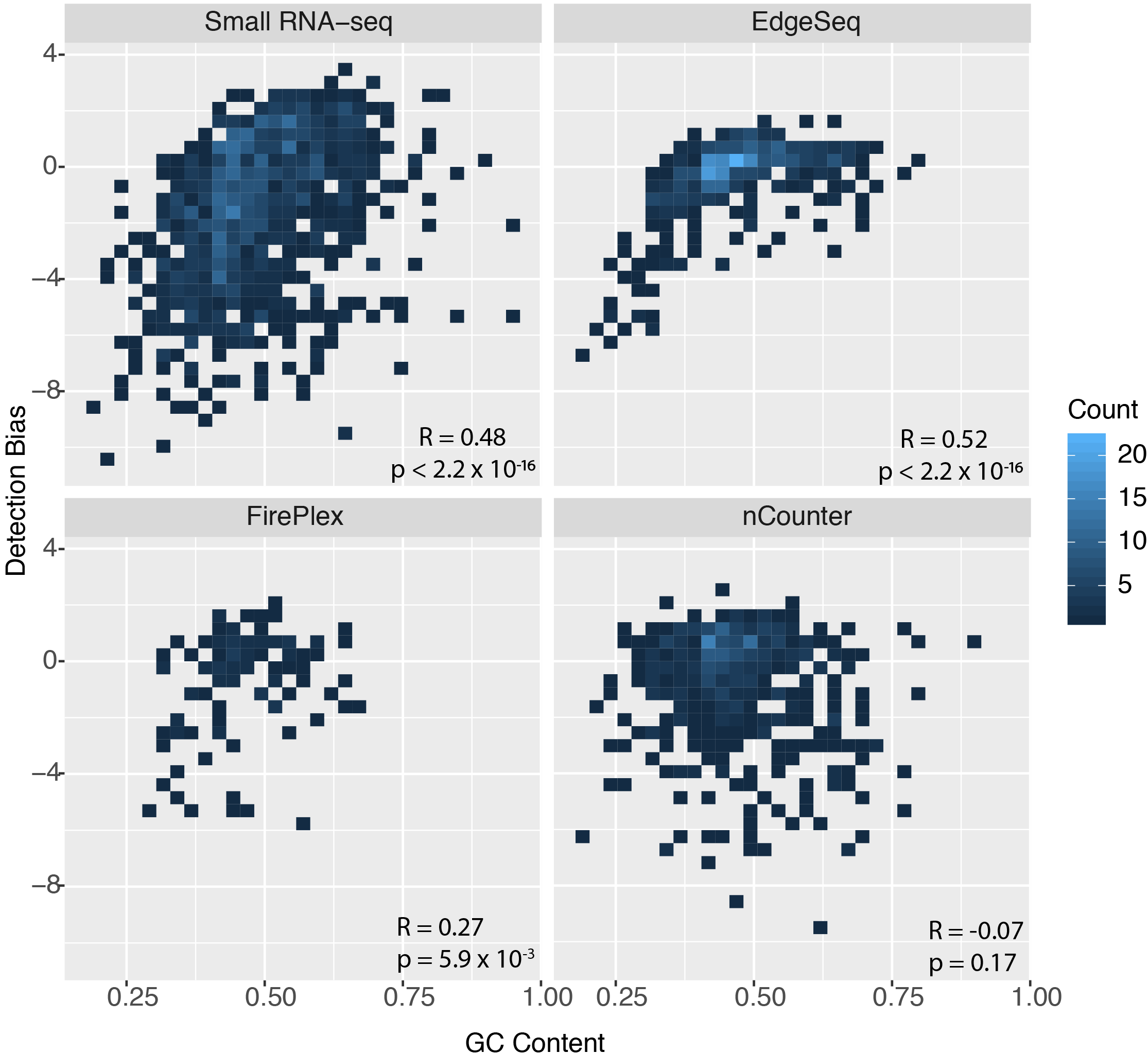
Relationships Between Bias and GC Content. Detection bias is plotted as a function of GC content. Lighter blue colors represent a higher density of miRNAs. Correlation coefficients and p-values were calculated using the Pearson method.

We next examined whether detection biases were consistent within platforms and compared biases across platforms. Biases were generally consistent within platforms, as demonstrated by correlating detection bias determined with one pool to detection bias determined with another pool (Supplemental Figure 3). As expected, biases were less well correlated between platforms (Fig. 4). The highest correlations were between EdgeSeq and small RNA-seq (R = 0.38) and EdgeSeq and FirePlex (R = 0.29). We conclude that detection biases for specific miRNA differ substantially between platforms.

**Figure 4.**
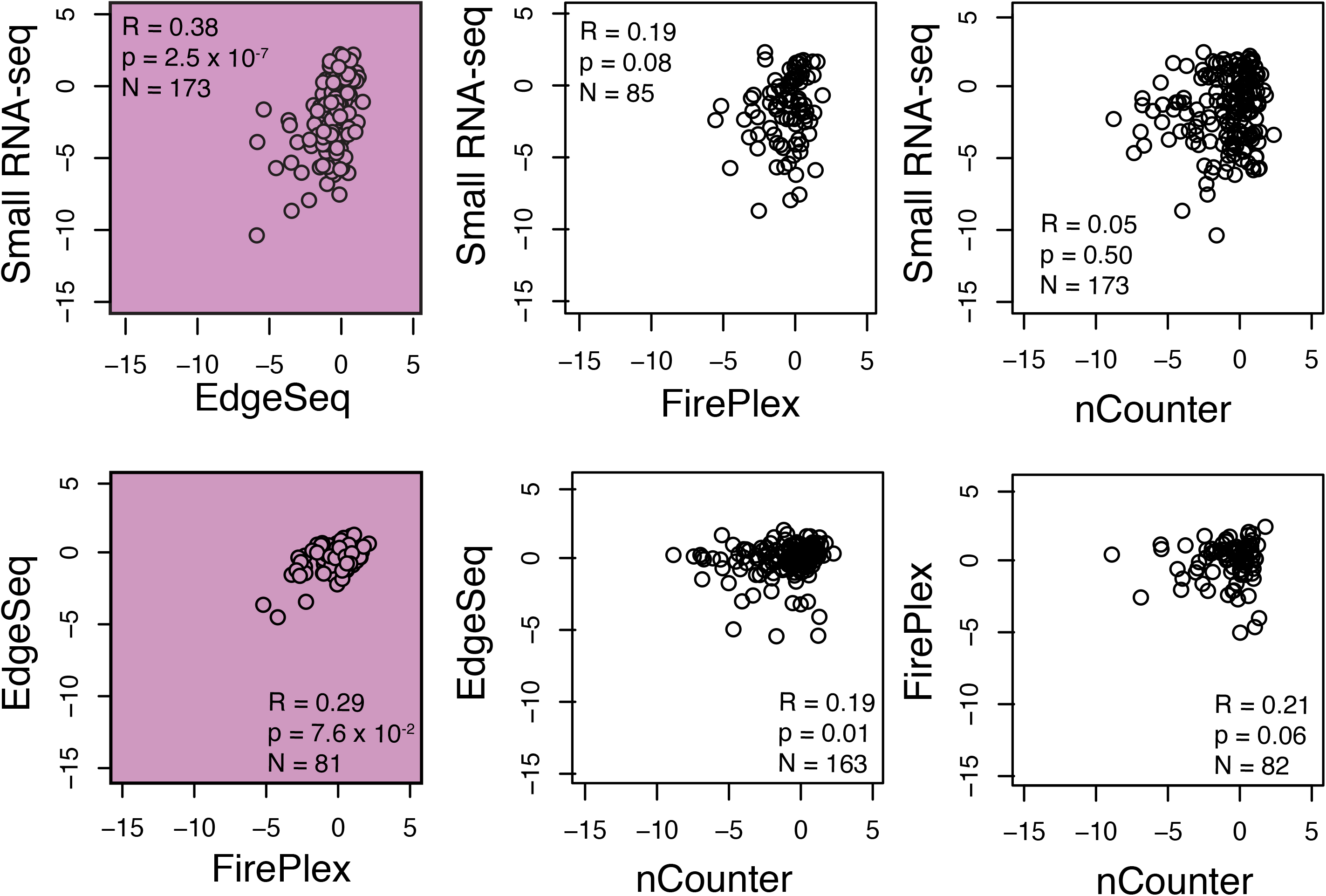
Detection Bias Comparison Across Platforms. Each point represents a pairwise comparison of the detection bias for each miRNA between the equimolar pool for the different platforms. Correlation coefficients and p-values are calculated using the Pearson method.

### Specificity and sensitivity analysis

We used data from the equimolar pool to determine the ability of each platform to distinguish between synthetic miRNAs that were present from those that were absent. Depending upon the platform, false positive signals for miRNAs not present in samples might be caused by a variety of phenomena, including incorrect nucleotide incorporation during reverse transcription or DNA amplification, sequencing errors, cross-hybridization to non-cognate miRNAs or other probes, or auto-fluorescence. All platforms showed some overlap between the distribution of signals for miRNAs that were present in the pool compared with those that were absent (Fig. 5A). As one means to assess false positives, we calculated the proportion of miRNAs that were absent from the pool but had signals higher than the fifth percentile of miRNAs that were present in the pool. This proportion was lowest for small RNA-seq (31/2081 miRNAs, 1.5%), intermediate for FirePlex (1/21, 4.8%), and nCounter (22/376, 5.9%), and highest for EdgeSeq (146/1632, 8.9%). By other metrics, separation between present and absent miRNAs was also best for small RNA-seq (ratio of median signal for present to median signal for absent = 1750; area under the receiver operating characteristic curve, AUC = 0.99), intermediate for EdgeSeq (ratio = 728, AUC = 0.97) and nCounter (ratio = 1078, AUC = 0.94), and least for FirePlex (ratio = 125, AUC = 0.81) (Fig. 5A, B). Overall, small RNA-seq was superior to the other platforms by each of these measures of sensitivity and specificity.

**Figure 5.**
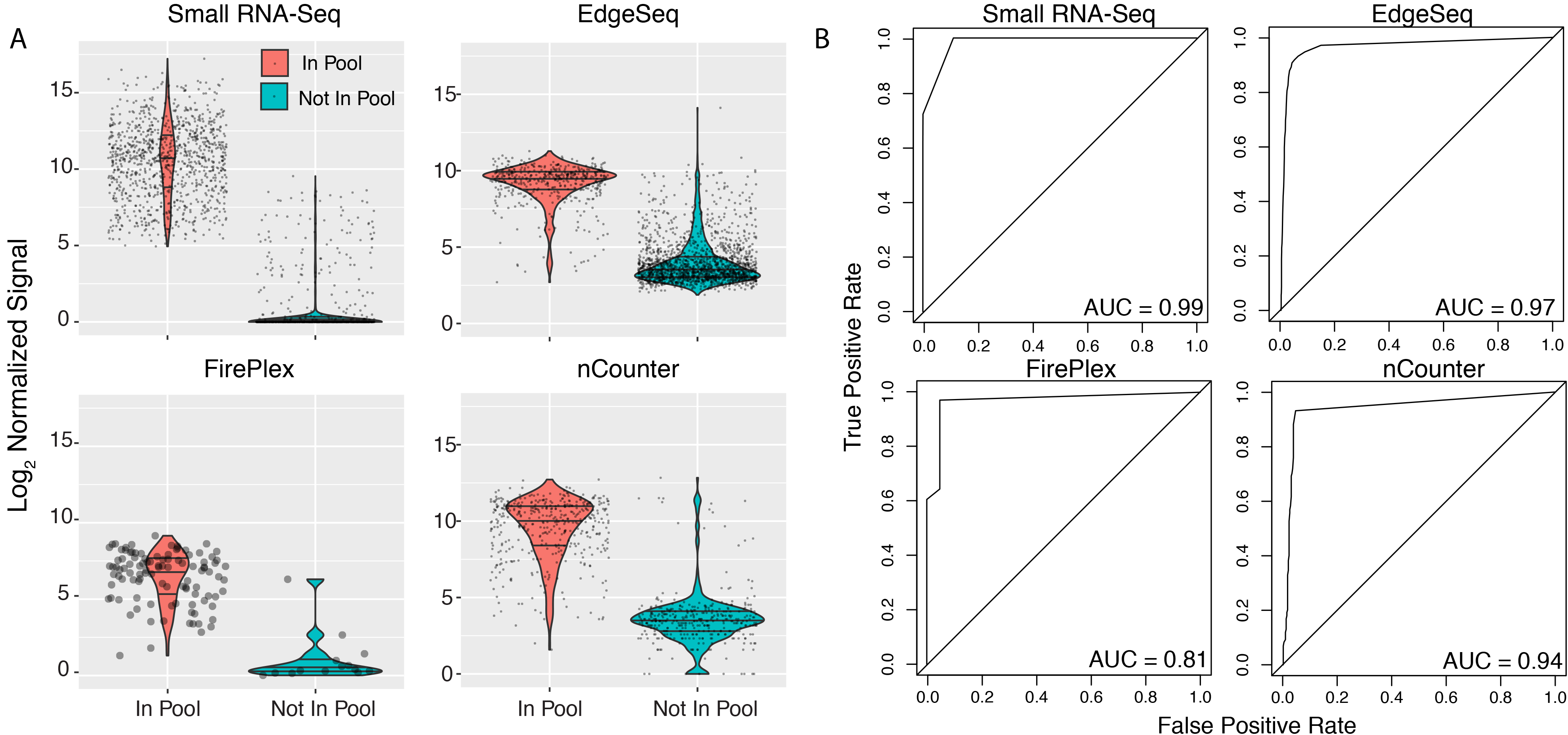
Sensitivity and Specificity of Each Platform As Assessed Using the Synthetic Equimolar Pool. (A) Normalized and log_2_-transformed signal is plotted for detectable miRNAs in the synthetic equimolar pool (small RNA-seq N = 758, EdgeSeq N = 451, FirePlex N = 50, nCounter N = 422) and those not in the synthetic equimolar pool (small RNA-seq N = 2,081, EdgeSeq N = 1,624, FirePlex N = 15, nCounter N = 376). (B) Receiver operating characteristic curves.

We investigated whether false positive signals could be related to cross-detection of miRNAs with similar sequences within the synthetic pool (Supplemental Figure 4). For EdgeSeq, the relatively small set of absent miRNAs with sequence similarity to present miRNAs did tend to have higher signals than other absent miRNAs. This was also observed with small RNA-seq but was not evident for nCounter. We were unable to assess FirePlex’s ability to distinguish closely related sequences since the smaller set of probes did not include any designed to recognize absent miRNAs that were similar in sequence to those present in the pool. Of the other three platforms, nCounter displayed the least evidence for cross-detection whereas EdgeSeq and to a lesser extent RNA-seq showed some evidence of cross-detection of closely related sequences.

### Relative Quantification of miRNAs

These four platforms are typically used for relative quantification – i.e., comparing the level of any particular miRNA between samples (e.g., case versus control). We used the ratiometric pools to assess each platform’s accuracy for relative quantification (Fig. 6). Each of the four platforms provided reasonably good estimates of ratios for most miRNAs, although all platforms were inaccurate for certain miRNAs. Root-mean-square-error (RMSE) for miRNA log ratios for RNA-seq (0.45), EdgeSeq (0.47), FirePlex (0.58), and nCounter (0.46) were quite similar. This evaluation with synthetic miRNA pools indicates that the four platforms behave similarly well for relative quantification.

**Figure 6.**
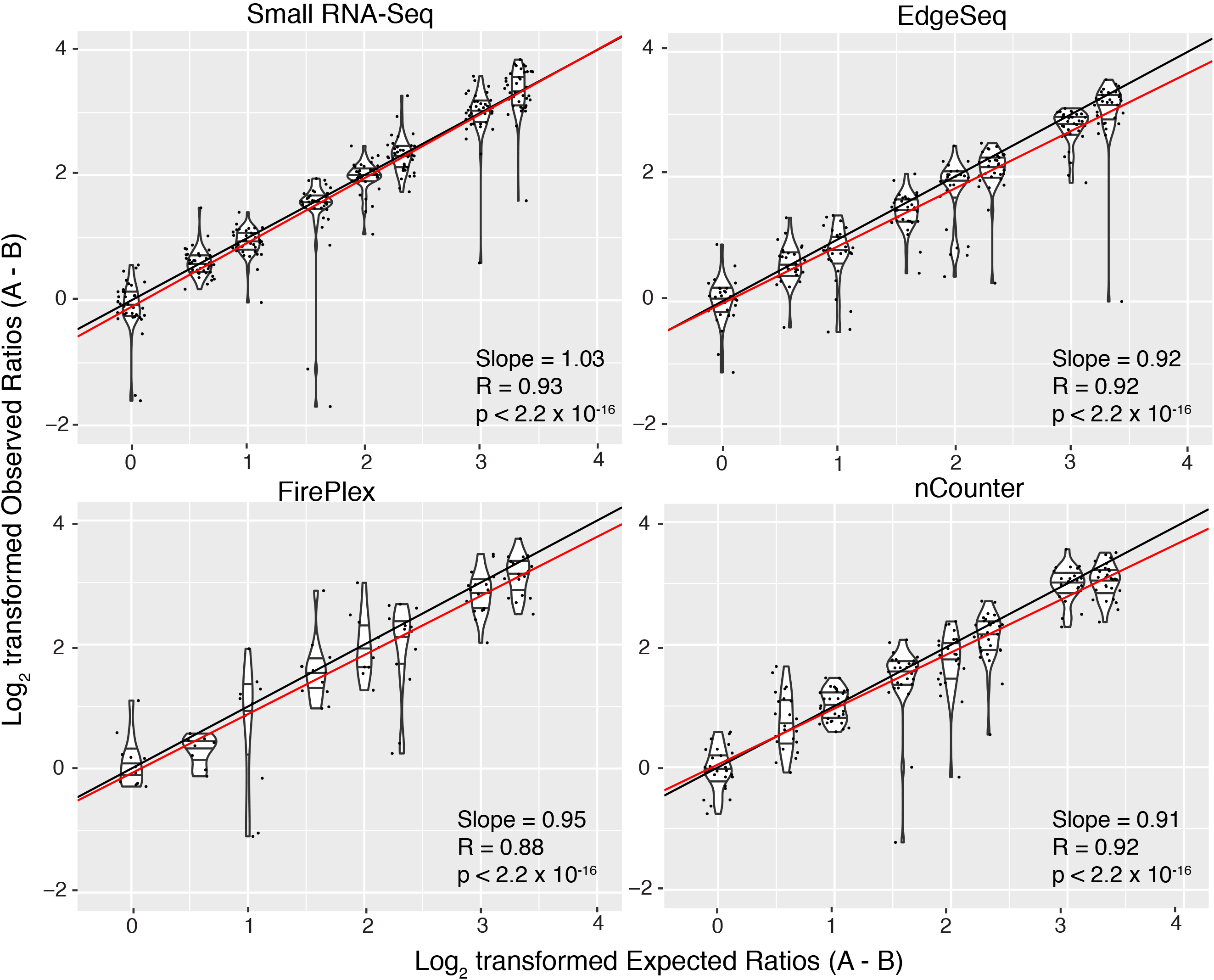
Accuracy of Relative Quantification Determined by Pools of Synthetic miRNAs at Ratiometric Concentrations. The expected fold-change of a miRNA in pool A and pool B is plotted against the observed fold-change, represented here as a log_2_ transformed ratio. Each violin plot represents the distribution of observed ratios for a particular expected ratio. The lines represent the median (middle) and the interquartile range (top and bottom). The dots represent the actual ratios. The black line is the line of identity, and the red line is the best fit line as calculated using linear regression.

### Analysis of reproducibility in plasma samples

To evaluate the reproducibility of each platform with biologically relevant samples, we analyzed exRNA isolated from human plasma samples in quadruplicate using small RNA-seq and in triplicate using EdgeSeq. We could not evaluate reproducibility of purified exRNA on FirePlex because RNA from these samples were not included in the FirePlex assay nor could we evaluate reproducibility of purified exRNA on nCounter because technical replicates were not performed by NanoString. Because EdgeSeq and FirePlex can also take as input a small volume of crude biofluid, we also analyzed exRNA directly from plasma in duplicate for EdgeSeq and triplicate for FirePlex. Reproducibility between technical replicates for RNA isolated from plasma was worse than that observed with the synthetic equimolar pool for small RNA-seq and EdgeSeq (Supplemental Figure 5, Table S3, Table S4), likely due to the lower average concentration of each miRNA in the plasma RNA samples compared to the synthetic pools. For isolated RNA, the median CV was higher with small RNA-sequencing (33.4%) than with EdgeSeq (14.4%). As previously reported for RNA-seq (Srinivasan et al., 2019), CVs decreased as signal increased, but overall CVs remained higher for the plasma RNA samples than for the synthetic pools.

For crude plasma, the median CV was lower with EdgeSeq (17.8%) than with FirePlex (43.2%) (Supplemental Figure 5). Signal intensities between data generated from isolated RNA versus crude plasma were moderately well correlated for EdgeSeq (R = 0.62, Supplemental Figure 6). To assess whether the use of crude biofluid biased against miRNAs carried in certain subcompartments in plasma, we inspected signals for miRNAs previously found to be differentially associated with 5 different subcompartments in human serum: CD63+ EVs, CD81/CD9+ EVs, AGO2+ RNPs, HDL, and lipoprotein-free fraction (LFF) (Srinivasan et al., 2019). We found no obvious systematic differences in signal according to the assigned subcompartment (Supplemental Figure 6), suggesting that EdgeSeq is capable of detecting miRNAs that are preferentially associated with each subcompartment within crude plasma samples.

### Analysis of miRNA in maternal plasma samples

We next compared miRNA signals across platforms using the mean normalized signal intensities of maternal plasma samples from two donors and observed moderate correlation between small RNA-seq and EdgeSeq (R = 0.68), small RNA-seq and FirePlex (0.78), and EdgeSeq and FirePlex (0.74) but weaker correlation with nCounter (small RNA-seq R = 0.43, EdgeSeq R = 0.53, FirePlex = 0.63) (Supplemental Figure 7). In the plasma samples, the weaker correlation between nCounter and the other three platforms appeared to be due to reduced sensitivity, as the nCounter measurements for the large majority of the miRNAs were near the lower limit of detection (median for negative control probes = 16.5, median for all human miRNA probes = 18). In addition, the differences in the patterns of bias between nCounter and the other methods (as seen with the synthetic pools in Figure 4) may also contribute to the weaker correlation.

To assess whether there was agreement in results among the different platforms when comparing biologically distinct samples, we examined the expression of a cluster of 50 placenta-specific miRNAs in chromosome 19 known to be specifically expressed during pregnancy (Ouyang et al., 2014). For all platforms, we compared signals for these miRNAs in maternal plasma samples versus a pool of non-maternal female plasma (see Methods). For all platforms, these miRNAs generally yielded signals that were near the lower end of the range for all miRNAs, suggesting that the placental miRNAs present in maternal plasma are generally less abundant than many other miRNAs found in both maternal and non-maternal plasma. A statistically significant increase in the expression of these miRNAs in the plasma of pregnant women compared to non-pregnant women was detected for 13/13 miRNAs in small RNA-seq (p = 1.2 × 10^−4^, Mann-Whitney test) (Fig. 7, Table S3, Table S4). EdgeSeq was also able to detect differences (10/11 miRNAs increased, p = 9.8 × 10^−4^), although fold-differences tended to be smaller than with RNA-seq. FirePlex (2/5 miRNAs increased, p = 0.41) and nCounter (4/12 miRNAs increased, p = 0.69) did not show a signficant difference in placental miRNAs. We conclude that RNA-seq and, to a lesser extent, EdgeSeq were able to detect differences in relatively low abundance miRNAs of placental origin, whereas FirePlex and nCounter were not.

**Figure 7.**
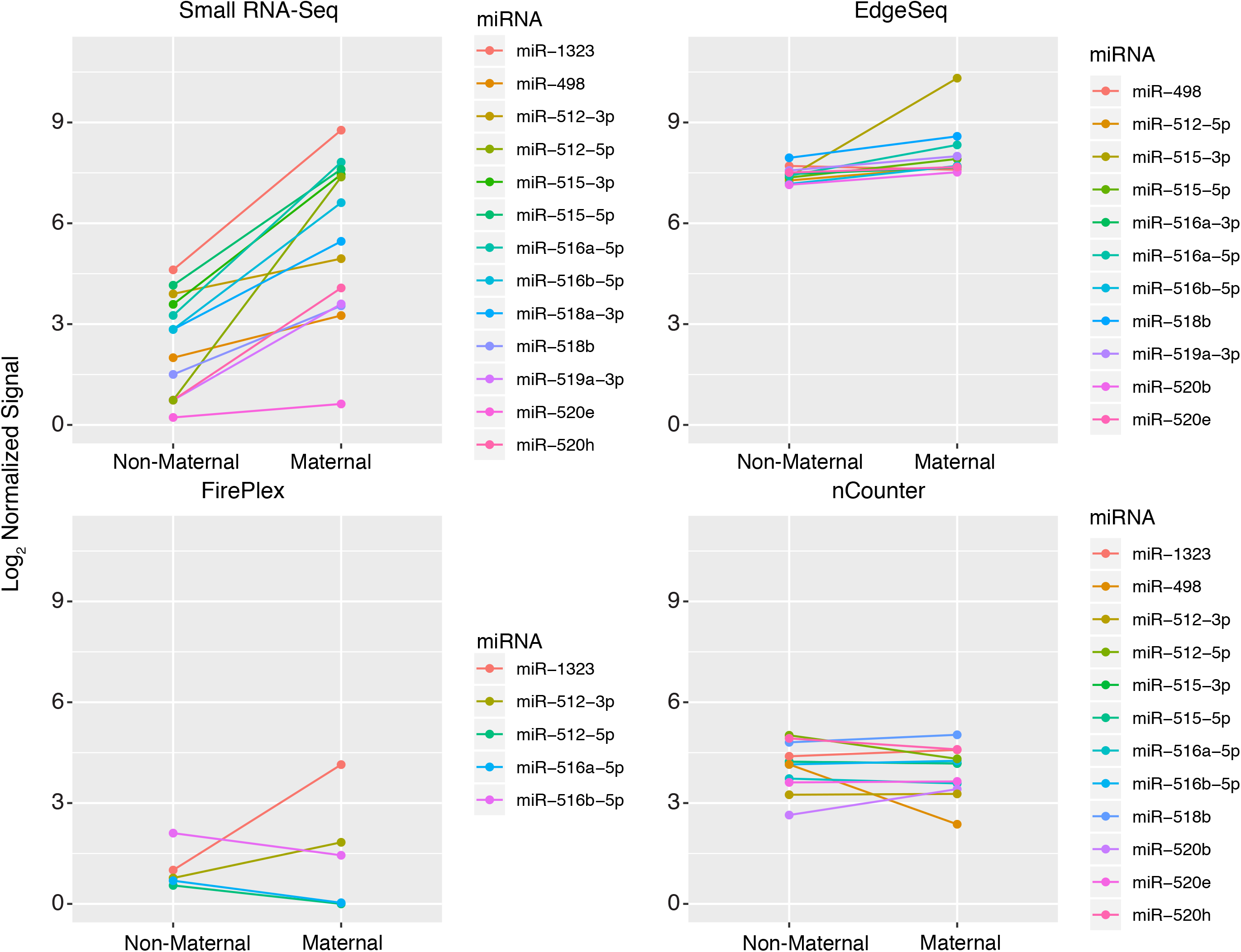
Expression of Maternal miRNAs in Non-Maternal Female Plasma and Maternal Plasma. Each pair of connected dots represents miRNA signals for non-maternal female plasma (pooled samples) and maternal plasma (mean of two maternal samples assayed separately).

## DISCUSSION

Two major objectives of Phase 1 of the ERCC were to identify methods for robust and reproducible quantification of extracellular miRNAs in biological fluids and to establish their utility as biomarkers. Here, we explored three miRNA measurement platforms as potential alternatives to small RNA-sequencing for relative quantification of extracellular miRNA. The platforms, which were utilized diverse technologies, were selected based on rapid turnaround time and ease of use, properties that are attractive for biomarker assays. The four platforms evaluated in this study included: a small RNA-sequencing protocol optimized for low input samples that had previously been compared to more widely-known commercial protocols (Giraldez et al., 2018); HTG Molecular’s EdgeSeq miRNA Whole Transcriptome Assay (EdgeSeq); Abcam’s FirePlex (FirePlex); and NanoString’s nCounter (nCounter). A recent report compared the performance of small RNA-seq, FirePlex, EdgeSeq, and Qiagen miRNome on standardized tissue samples and plasma exRNA samples (Yeri et al., 2018), but to our knowledge, a systematic study using synthetic RNA mixes that allow for accurate evaluation of assay performance has not been previously performed. Overall, our findings demonstrated that each platform had specific advantages and drawbacks that need to be taken into consideration when selecting a technology for exRNA studies, particularly those aimed at development and clinical application of extracellular miRNA biomarkers.

Our study evaluated the performance of each platform with respect to reproducibility, bias, specificity, and relative quantification using standardized samples. Our samples consisted of three pools of synthetic miRNAs, one of which contained the same concentration of each component miRNA (the equimolar pool) and two that contained different concentrations of the component miRNAs, designed such that the relative concentration of individual miRNAs between these two ratiometric pools ranged from 1:10 to 10:1. While these pools were critical for determining the bias, accuracy, and specificity of each platform, we also analyzed exRNA from pooled healthy male plasma, pooled healthy female plasma, and maternal plasma samples from two individual donors. These plasma exRNA samples allowed us to compare how the platforms performed on real biological samples.

Small RNA-sequencing is the only platform we studied that does not require hybridization, making it the only truly non-targeted platform. Given that small RNA-sequencing will capture any small RNA of acceptable (> 17 nt and < ~30 nt) length with a 5’ phosphate and 3’ hydroxyl group, it enables measurement of all RNA biotypes, which may be advantageous when studying clinically-useful biofluids in which miRNAs comprise a very small fraction of the exRNA, such as urine (Godoy et al., 2018). However, the turn-around time for small RNA-sequencing is slow, requiring at least a week for RNA isolation, cDNA library generation, sequencing, and analysis. Moreover, the reproducibility of small RNA-seq is strongly and negatively affected when different RNA isolation and library preparation protocols are used (Giraldez et al., 2018, Srinivasan et al., 2019). Even when the same protocols are used, reproducibility of measurements of plasma miRNAs with signal intensities less than 32 normalized counts was poor, potentially making it difficult to reliably quantify rare miRNAs across samples. Small RNA-seq is also impeded by bias introduced during adapter-ligation due to sequence-dependent T4 RNA ligase bias (Jayaprakash et al., 2011, Hafner et al., 2011). Although less biased than other commercially-available small RNA-sequencing kits (Giraldez et al., 2018), our small RNA-seq protocol exhibited higher bias compared to the three other tested platforms. This bias was partially dependent on GC content. Small RNA-sequencing detected a small number of reads that mapped to miRNAs not present in the synthetic equimolar pool, which may have been caused by errors introduced during PCR or sequencing or caused by contamination. Although most of these detected “not present” miRNAs had very low signal intensities, a few had greater than 500 normalized counts. Although all platforms could detect known differences in the synthetic ratiometric pools, small RNA-seq showed better detection of expected differences between biological samples that the other platforms, with the largest number of placenta-specific miRNAs having significantly higher levels of expression in maternal plasma compared to non-pregnant female plasma.

EdgeSeq is a high-throughput semi-automated assay that can process up to 96 samples in 24 hours, excluding the time for sequencing, with the multiplexing limit determined by the number of sequencing barcodes available. Additionally, its ability to use as little as 25 μL of crude plasma as input makes EdgeSeq an attractive platform for clinical use. Software for the alignment of fastq files and generation of miRNA counts is built into the same instrument that processes the samples, excluding the need for extra computational resources. For both crude plasma and isolated RNA technical replicates, EdgeSeq was more reproducible than small RNA-seq. One important advantage of EdgeSeq is its ability to directly assay miRNAs in small volumes of biofluid. Assays using crude plasma (25 μL) were only slightly less reproducible than those using RNA (isolated from 200 μL plasma). Comparisons of results from crude plasma and isolated RNA suggest that there are no major differences in the ability of EdgeSeq to detect miRNAs from these two different starting materials. On the other hand, the correlations between results from crude plasma and isolated RNA were not high enough to allow us recommend direct comparisons between data derived from these two input types. EdgeSeq did not suffer as strongly from detection bias as small RNA-seq; for the synthetic oligonucleotide pools, most miRNAs were detected at signal intensities near the expected signal. Although sources of bias in EdgeSeq are unknown, very low GC content was strongly associated with low signal intensity. EdgeSeq profiling of the equimolar pool showed high signal for several miRNAs not present in the equimolar pool, in extreme cases reaching signal intensities greater than the maximum signal intensity achieved by miRNAs in the pool. Although there was evidence of some signal from cross-hybridization, sequence similarity did not seem to explain most of the false positive signals. These signals may be attributable to other factors like incomplete digestion of unbound probes or cross-hybridization of probes in the assay. Placenta-specific miRNAs were detected at increased levels in maternal plasma than in non-maternal plasma by EdgeSeq, but fold-differences were modest compared to small RNA-seq, likely due to EdgeSeq’s lower sensitivity for quantification of low abundance miRNAs. This is consistent with small RNA-seq showing slightly better performance in relative quantification for the ratiometric pools (Figure 6). However, the true values of these miRNAs are unknown, and so the accuracy of the fold-change measurements could be ascertained.

Like EdgeSeq, FirePlex can use as input a small volume (20 μL) of crude plasma. The number of miRNAs that can be analyzed in a single FirePlex assay is smaller than for the other methods, but FirePlex miRNA panels are easily customized, which offers the advantage of only focusing analysis on miRNAs of interest. Processing of samples takes approximately 1.5 days and requires the use of a flow cytometer for signal detection. Of all tested platforms, FirePlex was the least reproducible for the synthetic samples and had similar reproducibility as small RNA-seq for the plasma samples. FirePlex had low bias, with median signal intensities close to the expected signal. The source of detection bias in FirePlex is unknown and was only slightly affected by GC content. A small number of miRNAs not present in the synthetic equimolar pool were detected at levels comparable to those present in the pool, but because there were no FirePlex probes for sequences within an edit distance of 4 from these false positives, we could not determine whether sequence similarity contributed to false positives. Although we evaluated reproducibility of technical replicates of crude plasma, we did not perform the FirePlex assay using RNA isolated from the same crude plasma sample, so could not compare the reproducibility of crude plasma vs purified exRNA for this platform. For FirePlex, all of the placenta-specific miRNAs were detected with low signal, and thus the failure to detect differences may be due to poor reproducibility of measurements near the threshold of detection.

While nCounter does not accept crude plasma as input, it can assay up to 800 miRNAs, making it more a higher-throughput assay than FirePlex. The turn-around time for the nCounter assay is approximately 2 days. Like EdgeSeq, software for fluorescence detection, demultiplexing, background-subtraction, and reporting of signal intensities, is incorporated into the processor. One limitation to our study is that technical replicates were not performed by NanoString so reproducibility could not be assessed for nCounter. nCounter had less bias than RNA-seq but more bias than the other two platforms. GC content did not seem to have any effect on the detection bias, and the detection bias for nCounter was not correlated with any other platform, suggesting unique biases. nCounter involves direct detection and eliminates potential bias associated with amplification. However, our analysis indicates that the other aspects of the assay are subject to bias and that results from nCounter and the other three platforms tested cannot be used to directly infer absolute miRNA quantities. We observed that for all of the plasma exRNA samples, nCounter yielded a much smaller fraction of miRNAs detected above background compared to the other three platforms, and thus the failure to detect differences between pregnant and non-pregnant serum could be due to lower sensitivity of this platform.

Our study reveals differences in performance that are relevant when deciding on a platform for a specifc application. Small RNA-seq is well suited for discovery applications, had the best ability to distinguish between miRNAs that were present in or absent from the synthetic pool, and was best able to meet the demanding challenge of detecting placental miRNAs in maternal plasma. However, due to throughput and turnaround time considerations, small RNA-Seq may not be practical for clinical use. EdgeSeq offers the ability to detect a large number of miRNAs in either isolated RNA samples or crude plasma, had relatively low detection bias, and had some success in detecting placental miRNAs in maternal plasma. FirePlex, with a smaller number of probes per assay, is best suited for more targeted analyses and like EdgeSeq can be used with crude plasma as well as isolated RNA, but this platform did not offer clear advantages in terms of reproducibility, bias, sensitivity and specificity, or relative quantification compared with the other methods and could not detect placental miRNAs in maternal plasma. For nCounter, we were unable to assess reproducibility, but we found that the absence of an amplification step did not reduce bias below that seen with the other two probe-based platforms and we were unable to detect placental miRNAs in maternal plasma. With the exception of RNA-seq, our observations are based on results provided to us by manufacturers of these platforms, and subsequent versions of these platforms may have different performance characteristics. For example, we observed that the version of the FirePlex assay analyzed here was superior for relative quantification but was less reproducible than an earlier version tested with some of the same samples (data not shown). The analyses that we report here provide a useful comparison of available platforms for miRNA quantification and help highlight limitations that could be addressed in future technologies. In the meantime, platform-specific issues related to reproducibility, bias, sensitivity, and specificity must be taken into account when interpreting the results of extracellular miRNA analyses in biomarker and other studies.

## EXPERIMENTAL PROCEDURES

### Synthetic RNA pools

The three pools of synthetic miRNAs were distributed to members of the NIH Extracellular RNA Communication Consortium (ERCC) and have been described previously (Giraldez et al. 2018). Briefly, the equimolar pool was produced by combining 962 human and non-human synthetic miRNAs from the miRXplore Universal Reference (Miltenyi Biotech) with a custom set of 190 other RNA oligonucleotides at equal molar concentrations (total of 1,152 RNA oligonucleotides, including 759 that correspond to known human miRNA sequences). The ratiometric pools contain 334 synthetic miRNAs, 286 of which correspond to human miRNA sequences, and were generated by IDT.

### Plasma samples

Human male and female plasma samples from healthy donors 21-45 years of age were used to assess reproducibility and maternal miRNA expression for small RNA-seq, EdgeSeq, and FirePlex. These were collected, processed, and distributed by the laboratory of Dr. Ionita Ghiran at Beth Israel Deaconess Medical Center (BIDMC). Details regarding IRB approval and collection of plasma are in Giraldez et al. (2018).

Non-maternal and maternal plasma was collected from donors ≥ 18 years of age under an IRB protocol approved by the Human Research Protections Programs at the University of California San Diego (UCSD). Biofluid samples were labeled with study identifiers; no personally identifiable information was shared among participating laboratories. Non-maternal female plasma was collected from 10 healthy non-pregnant adult donors, 22-26 years of age. Briefly, for each donor, ~500mL of whole blood was exsanguinated using 19 gauge needles (MONOJECT ANGEL WING Blood Collection Set 19G × 3/4,” ITEM# 79027, MFG#8881225281, Moore Medical) and 60 mL syringes, with 440 ul 0.5M K2EDTA added, from a peripheral vein and transferred to 50 mL conical tubes. The uncoagulated blood was then centrifuged at 2000 x g for 20 min to remove cells and cell debris. The clear supernatant was transferred to fresh tubes. The resulting cell-free plasma were pooled using equal volumes from each donor. These pools were split into 1.5 mL aliquots and stored at −80° until analyses were performed.

Maternal female plasma was collected from two healthy pregnant donors (36-37 years of age) during the 2^nd^ and 3^rd^ trimester. Briefly, for each donor approximately 8mL of whole blood was collected by peripheral venipuncture into a BD Vacutainer K2 EDTA plasma collection tube (Becton Dickinson PN 366643) at room temperature and centrifuged at 2000 x g for 20 minutes. The upper cell free plasma fraction was divided into 1 mL aliquots and stored at −80°C until analyses were performed.

### Small RNA-seq, EdgeSeq, FirePlex, and nCounter assays

For small RNA-seq of the synthetic equimolar and ratiometric pools, a total molar concentration of 10 femtomoles were used as the starting input. Total RNA from the pool of healthy human male plasma (for reproducibility analysis) was isolated as described in Giraldez et al. (2018); 2.1 μL of isolated RNA was used as input, corresponding to ~84 μL of plasma. Total RNA from the pool of non-maternal female plasma and the two maternal plasma samples (for maternal miRNA expression analysis) was isolated using a modified version of the QIAGEN miRNEasy Micro Kit as described in Godoy et al. (2018); 5 μL of isolated RNA was used as input, corresponding to ~200 μL of plasma. RNA from the pool of healthy human male plasma was run in quadruplicate; RNA from all female plasma samples was run in duplicate. All sequencing libraries were generated using the 4N protocol D as previously described (Giraldez et al., 2018). Synthetic equimolar pools were sequenced to a median read depth of 12,569,524 for total reads and 11,230,216 for miRNA reads, 11,130,839 and 5,987,818 for synthetic ratiometric pools, 23,262,217 and 2,367,895 for human male plasma, 17,823,762 and 2,072,402 for human female plasma, and 11,901,781 and 3,314,000 for maternal plasma samples. The libraries corresponding to the synthetic pools and the pool of healthy human male plasma were sent to the laboratory of Muneesh Tewari at the University of Michigan, Ann Arbor for sequencing on an Illumina HiSeq 2500. For all other plasma samples, sequencing was done at the Center for Advanced Technology at the University of California, San Francisco on a HiSeq 2500. Fastq files were shared and aligned via the extracellular RNA processing toolkit (exceRpt) pipeline (Rozowsky et al., 2019). Sequences smaller than 17 nt were removed prior to alignment. No mismatches were allowed when aligning to the synthetic pools; 1 mismatch was allowed for the human plasma libraries. Small RNA-seq results for the synthetic pools and pools of healthy human male plasma have been previously reported (Giraldez et al., 2018).

We shipped 40 nM stocks of the synthetic equimolar and synthetic ratiometric pool to HTG Molecular that were then diluted to 1 pM in HTG Lysis Buffer just prior to processing. 25 μL of the dilution was added to each well for a total molar concentration of 0.025 femtomoles. Crude plasma from the pool of healthy human male plasma, pool of healthy human female plasma, and two maternal plasma samples was prepared by adding 40 μL of sample to 40 μL of HTG Plasma Lysis Buffer followed by addition of 8 μL of Proteinase K, and incubated for 180 minutes. 25 μL were used as the final volume for each replicate, corresponding to 13 μL of plasma. For RNA from the pool of healthy human male plasma, 5 μL of RNA (corresponding to ~200 μL of plasma) isolated as described in Godoy et al. (2018) were added to 20 μL of HTG Lysis Buffer per reaction. For all reactions, the final volume was 25 μL. All samples were run on the HTG EdgeSeq Processor using the HTG EdgeSeq miRNA WT assay, which included 2,102 total probes, 2,083 of which were designed to recognize human miRNAs. Three technical replicates were run for each of the three synthetic samples, the pool of human male plasma, and RNA isolated from the human male plasma. Nine technical replicates of maternal plasma and non-maternal female plasma were run. Barcodes and adapters were added to processed samples using 16 cycles of PCR. Libraries were sequenced on an Illumina MiSeq.

We shipped RNA isolated from the human male plasma and crude maternal plasma from 2 donors, along with the synthetic pools and pools of human male and female plasma to Abcam for processing with the FirePlex assay. This assay included 131 total probes, all of which were designed to recognize human miRNAs. miRNAs were included in the panel based on their presence in the synthetic pools and plasma small RNA-sequencing libraries and covered a broad range of expression. 5 of the placental-specific miRNAs were also included in the panel. Two separate panels were run in order to accommodate all 131 total probes, as FirePlex is limited to 68 fluorescent barcodes per panel. The first panel contained 66 miRNA probes, the second contained 65. 6 miRNAs were present in both panels; for these miRNAs only the signal intensity for the miRNA in the first panel was kept. For each panel, 0.57 femtomoles of the synthetic equimolar pool was added to each well. For the ratiometric pool, 0.33 femtomoles were added. 20 μL of crude plasma was used for each replicate. All samples were run in triplicate.

The nCounter assay included 828 total probes, 798 of which were designed to recognize human miRNAs. The target concentration for each miRNA in the synthetic pools was 10 attamoles. Therefore, for the equimolar synthetic pool, which contained ~1200 RNA sequences, we sent the manufacturer 3 picomoles total at a concentration of 12 femtomoles/μL. For each ratiometric pool, which contained 334 RNA sequences, 1.5 picomoles total at a concentration of 3.4 femtomoles/μL were sent. For the exRNA samples isolated from maternal and non-maternal plasma using the RNA isolation method described in Godoy et al. (2018), we sent the equivalent of the amount contained in 20 μL of plasma, as recommended by the manufacturer. The manufacturer (Nanostring) analyzed the samples on the nCounter Human v3 miRNA panel.

### Detectable miRNA criteria

For small RNA-sequencing, we mapped reads to the complete set of synthetic RNA oligonucleotides in each pool. The equimolar pool contained 164 synthetic RNAs that were outside of the miRNA size range (<18 nt or >30 nt) or were not 5’ phosphorylated and these were excluded from the analysis. Only 759 of the 988 detectable miRNAs were human miRNAs; however, non-human miRNAs are still considered detectable for small RNA-seq as they are the appropriate length and contain the appropriate end modifications. Non-human miRNAs consisted of rat, mouse, and viral miRNAs. For the other platforms, detectable miRNAs were defined as those which were targeted by a probe in the assay.

### Signal Intensity

For small RNA-seq and EdgeSeq, signal intensities were determined from the numbers of mapped reads. For FirePlex and nCounter, the manufacturer adjusted the signal intensities by subtracting background fluorescence from raw signal intensities. For all analyses, signal intensities were quantile normalized using the normalizeQuantile function in the R library limma (Ritchie et al., 2015).

### Analysis of reproducibility

To assess reproducibility of the synthetic miRNA pools and the human plasma samples, coefficient of variation (CV) was calculated as a percentage using the cv function in the R library raster (Hijmans 2019). For analysis of the relationship between CV and signal intensity, miRNAs were then divided into 5 equal-sized groups for the synthetic pools and 4 equal-sized groups for the plasma samples according to median signal intensity across replicates. All plots were produced using ggplot2 (Wickham 2009).

### Analysis of Bias

To analyze bias, we calculated bias as the ratio of observed signal to expected signal. For the equimolar pool, the expected signal was defined as the total signal intensity (sum of averaged normalized signal intensities across technical replicates for all detectable miRNAs) divided by the number of detectable miRNAs. For the ratiometric pools, the expected value for any given miRNA was made proportional to the relative concentration of that miRNA (e.g. a miRNA at 10X had a signal intensity 10 times greater than a miRNA at 1X). We used the Spearman method (cor function in R library stats) to correlate GC content and detection bias. To evaluate whether there was a relationship between the 5’ or 3’ nucleotide and normalized signal, we performed Mann-Whitney tests between each combination of nucleotides using the wilcox.test function in the R library stats. P-values were corrected using the Bonferroni method using the p.adjust function in the R library stats (R Core Team 2018). Correlation of bias in the synthetic equimolar pools (observed/expected) across platforms was calculated using the Pearson method.

### Analysis of Sensitivity and Specificity

For small RNA-seq, “absent miRNAs” were defined as all miRNAs in miRbase (version 21) that were not present in the synthetic pool. We only considered miRNAs in the synthetic pools that were human. For EdgeSeq, FirePlex, and nCounter, “absent miRNAs” were defined as all miRNAs that had probes but were not present in the synthetic pool.

To generate receiver operator characteristic curves, the true positive rate and false positive rate were calculated using only the signal intensities from the synthetic equimolar pool for each platform. We set each threshold as the expected value for each platform multiplied by all numbers from 0 to 200 with an increment of 0.05. The number of true positives was calculated as the number of miRNAs in the synthetic equimolar pool with a signal intensity greater than or equal to the threshold. Similarly, the number of false negatives was equal to the number of miRNAs with a signal intensity less than the threshold. In the same manner, the number of false positives and true negatives were calculated for miRNAs not in the synthetic equimolar pool. The true positive rate is calculated as the number of true positives divided by the sum of the number of true positives and false negatives. The false positive rate is calculated as the number of false positives divided by the sum of the number of false positives and true negatives. The area under the curve was calculated using the AUC function in the R library DescTools (Signorell et al., 2019).

We calculated the minimum Levenshtein edit distance between each absent miRNA and any miRNA present in the pool using the Levenshtein python package.

### Analysis of Relative Quantification using the Synthetic Ratiometric Pools

The expected count for each miRNA was calculated by dividing the sum of all miRNA signal intensities by the relative number of detectable miRNAs. For example, in EdgeSeq, there were 242 overlapping miRNAs between the ratiometric pool and the EdgeSeq probe set. However, because miRNAs were present at varying concentrations (e.g. 139 at 1X, 14 at 1.5X, 14 at 2X), the sum of signal intensities was not divided by 242 but by the sum of the number of miRNAs multiplied by the relative concentration (e.g. 139*1 + 14 * 1.5 + 14*2 + …). Once the expected count was calculated, it was adjusted for each miRNA based on its relative concentration.

The expected ratio for each miRNA was calculated as the concentration of a miRNA in pool A divided by the concentration in pool B, unless the concentration in B was higher than in A, in which case the expected ratio was the inverse. The observed ratio for each miRNA was calculated from quantile normalized intensities.

The best fit line was calculated using the lm function in the R library stats, and the correlation coefficient and p-value were calculated using the Pearson method using the cor function in the R library stats.

### Analysis of Maternal miRNA Expression in Human Plasma Samples

The 50 miRNAs arising from chr19:53666679-53788773 (hg19) were considered to be maternal-specific miRNAs (Ouyang et al., 2014). Technical and biological replicates were quantile normalized and log_2_-transformed signal intensities from maternal and non-maternal samples were compared for all of the 50 miRNAs that were detected in each platform. Since 9 technical replicates were run for the non-maternal female plasma and maternal plasma, we randomly chose 3 replicates for this analysis. For small RNA-seq, EdgeSeq, and nCounter, the non-maternal female comparator was the pool of healthy female plasma collected at UCSD. For Fireplex, the comparator was the pool of healthy female plasma collected at BIDMC.

### Accession Numbers

## AUTHOR CONTRIBUTIONS

P.M.G., A.J.B, S.D., D.J.E., and L.C.L. designed the study. P.M.G. isolated RNA from all plasma samples, generated cDNA libraries, prepared samples for sequencing. D.J.E., A.J.B., and P.M.G. handled all correspondence with HTG Molecular. S.D. was the main correspondent with Abcam. L.C.L., P.D., and S.S. worked with NanoString to process samples on the nCounter. P.M.G. analyzed the data. P.M.G., D.J.E., S.D., and L.C.L. wrote the paper. All authors contributed to manuscript editing.

## ACKNOWLEDGMENTS

This publication is part of the NIH Extracellular RNA Communication Consortium paper package and was supported by the NIH Common Fund’s exRNA Communication Program. We would like to thank Ionita Ghiran for the pools of healthy human male and female plasma, and Aileen Fernando, Fabian Flores, and the UCSD Clinical and Translational Research Institute for assisting with collection of the maternal and non-maternal female samples used in this study. We would like to thank Prescott Woodruff who is a co-principal investigator on the grant supporting this study and William Thistlethwaite for help on data deposition. This work was supported by the NIH Extracellular RNA Communication Program (grant U01HL126493 to P.G.W. and D.J.E., UH3TR000906 and U01HL126494 to L.C.L., and UH3TR000901 to S.D.).

## SUPPLEMENTAL INFORMATION

The following tables are provided separately as Supplementary Data files in Excel format.

Table S1. Normalized Read Counts of Synthetic Equimolar Pool from Small RNA-sequencing. Related to Figures 1-5.

Table S2. Normalized Read Counts of Synthetic Equimolar and Ratiometric Pool from EdgeSeq, FirePlex, and nCounter. Related to Figures 1-6.

Table S3. Normalized Read Counts of Synthetic Ratiometric Pool from Small RNA-sequencing. Related to Figures 2,6.

Table S4. Normalized Read Counts of Plasma Samples from Small RNA-sequencing, EdgeSeq, FirePlex, and nCounter. Related to Figure 7.

