## Supplemental Figures for "Comparison of miRNA profiling methods using synthetic miRNA pools and standardized exRNA samples reveals substantial performance differences"

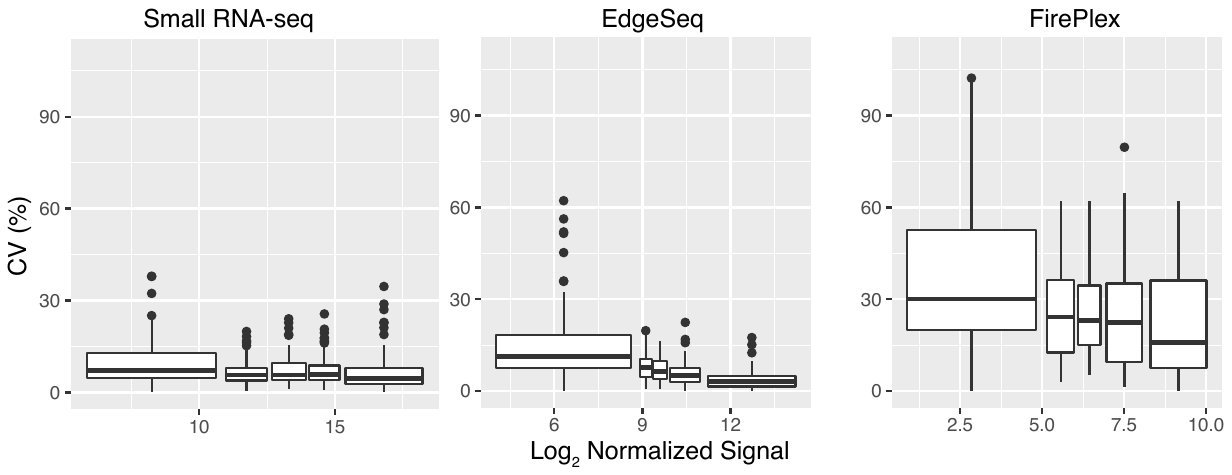

**Supplemental Figure 1. Reproducibility of small RNA-seq, EdgeSeq, and FirePlex when using Synthetic Ratiometric Pools.**

Coefficient of variation for technical replicates (small RNA-seq N = 4, EdgeSeq N = 3, FirePlex N = 3) expressed as a percentage as a function of median signal. Each boxplot represents ~20% of the total number of detectable miRNAs in the synthetic ratiometric pools, grouped by ascending expression (for total number of detectable miRNAs: small RNA-seq N = 296, EdgeSeq N = 242, FirePlex N = 100). Boxes represent median and interquartile ranges, whiskers represent 1.5 times the interquartile range. Dots represent outliers.

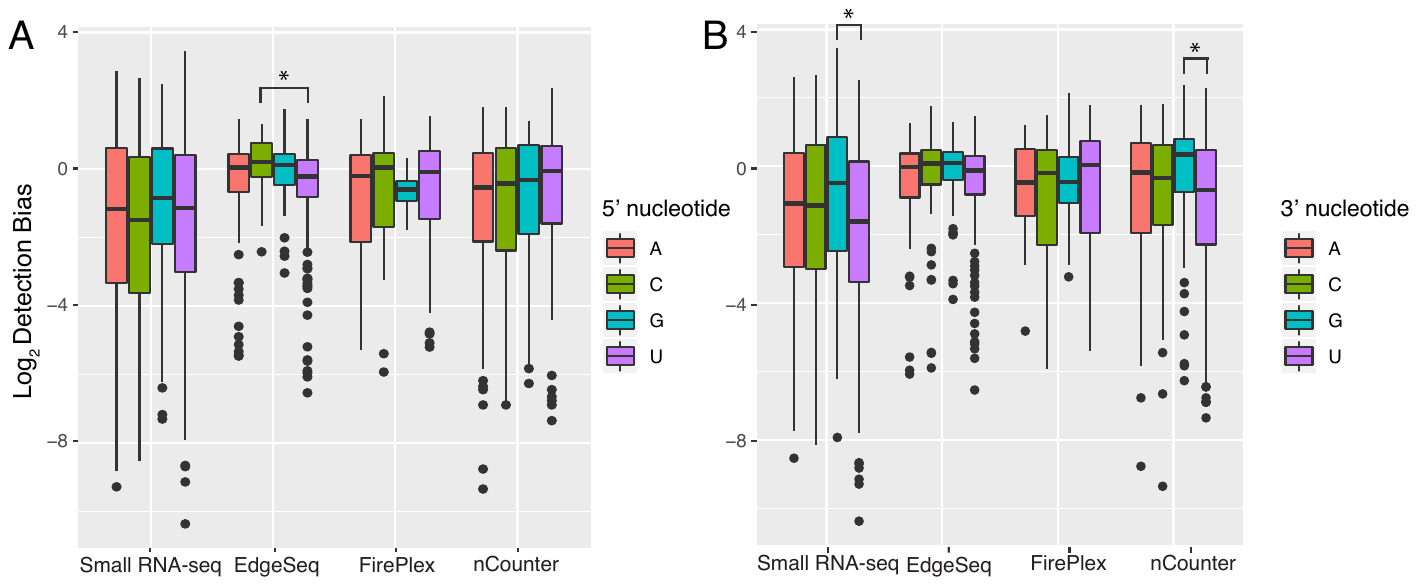

**Supplemental Figure 2. Relationship between bias and 5’ and 3’ nucleotide.**

Detection bias (ratio of observed to expected signal) is plotted grouped by either the 5’ (A) or 3’ (B) nucleotide. Boxes represent median and interquartile ranges, whiskers represent 1.5 times the interquartile range. Dots represent outliers.

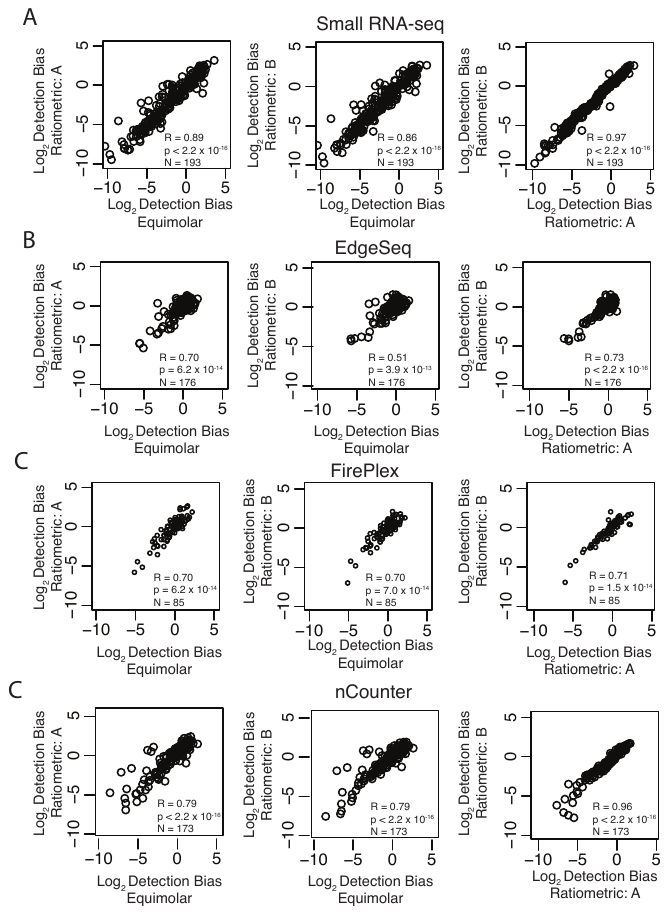

**Supplemental Figure 3: Comparison of detection bias across different synthetic pools of miRNAs.**

Each point represents a single miRNA that was present in the pools. Correlation coefficients and p-values were calculated using the Pearson method.

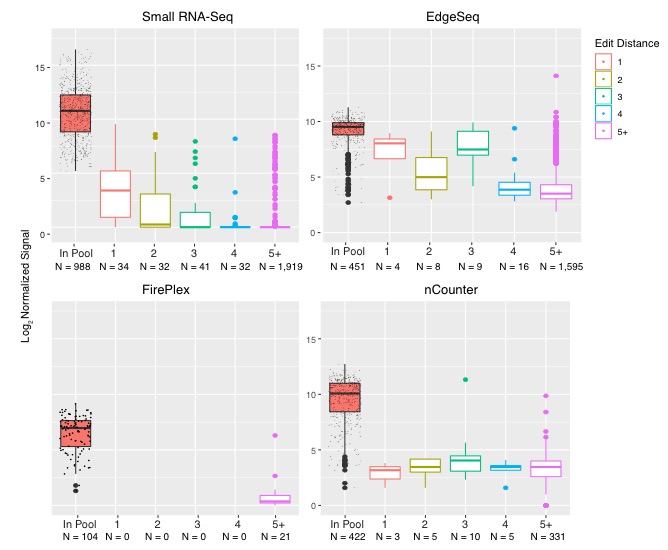

**Supplemental Figure 4**: **Specificity as assessed by the synthetic equimolar pool.**

Red violin plots represents the distribution of reads from miRNAs in the equimolar pool. All other violin plots represent the distribution of reads from miRNAs not in the equimolar pool with a Levenshtein edit distance of 1, 2, 3, 4, or more.

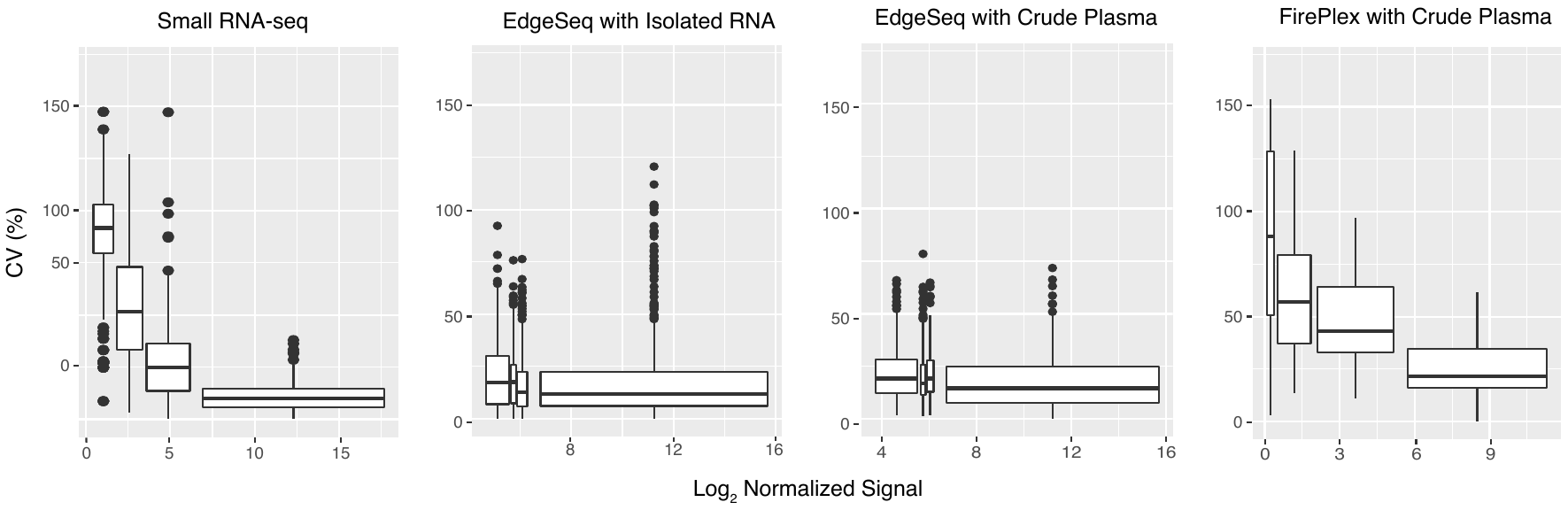

**Supplemental Figure 5. Reproducibility of small RNA-seq, EdgeSeq, and FirePlex when using Human Plasma Samples.**

Coefficient of variation for technical replicates (small RNA-seq N = 4, EdgeSeq N = 3, FirePlex N = 3) expressed as a percentage as a function of median signal. Each boxplot represents ~25% of the total number of miRNAs detected in the pool of healthy human male plasma, grouped by ascending expression (for total number of detectable miRNAs: small RNA-seq N = 1290, EdgeSeq N = 2083, FirePlex N = 125). Boxes represent median and interquartile ranges, whiskers represent 1.5 times the interquartile range. Dots represent outliers.

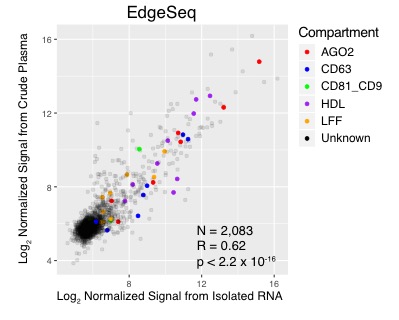

**Supplemental Figure 6. Comparison of miRNA Signals when using Isolated RNA or Crude Plasma.** Each point represents a pairwise comparison of the log_2_-transformed normalized signal from miRNA obtained from either a starting input of isolated RNA or crude plasma. Colored dots represent miRNAs with an associated carrier subclass. Dots are colored black and transparent if the subcompartment of the miRNA is unknown.

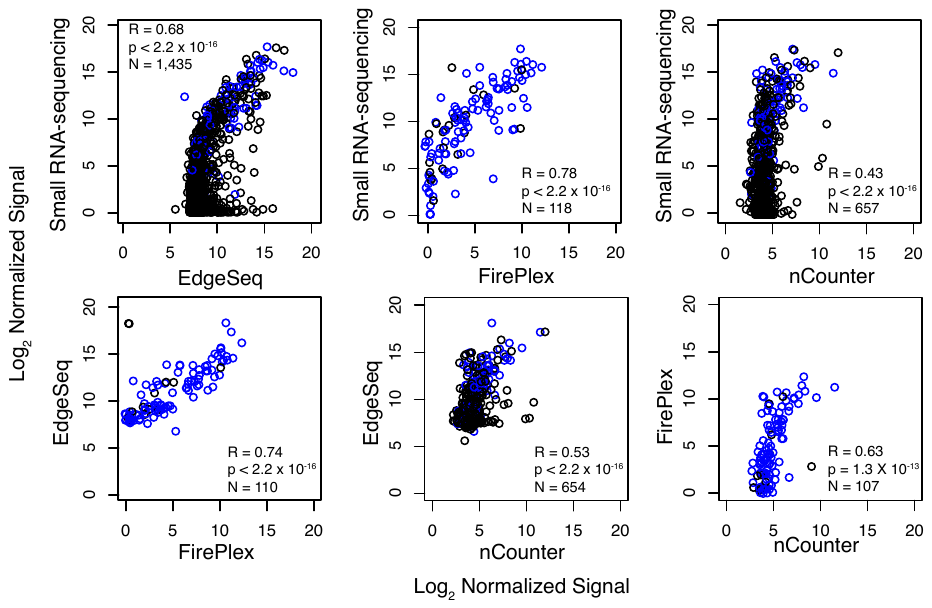

**Supplemental Figure 7. Pairwise Comparison of miRNA Signals from Maternal Plasma.** Each point represents a pairwise comparison of miRNA signal from each of the four platforms. Black points represent miRNAs in common between the pairwise comparison; blue points represent miRNAs in common across all platforms.
